# Tumor-localized interleukin-2 and interleukin-12 combine with radiation therapy to safely potentiate regression of advanced malignant melanoma in pet dogs

**DOI:** 10.1101/2024.02.12.579965

**Authors:** Jordan A. Stinson, Matheus Moreno P. Barbosa, Allison Sheen, Noor Momin, Elizabeth Fink, Jordan Hampel, Kimberly Selting, Rebecca Kamerer, Keith L. Bailey, K. Dane Wittrup, Timothy M. Fan

## Abstract

The clinical use of interleukin-2 and -12 cytokines against cancer is limited by their narrow therapeutic windows due to on-target, off-tumor activation of immune cells when delivered systemically. Engineering IL-2 and IL-12 to bind to extracellular matrix collagen allows these cytokines to be retained within tumors after intralesional injection, overcoming these clinical safety challenges. While this approach has potentiated responses in syngeneic mouse tumors without toxicity, the complex tumor-immune interactions in human cancers are difficult to recapitulate in mouse models of cancer. This has driven an increased role for comparative oncology clinical trials in companion (pet) dogs with spontaneous cancers that feature analogous tumor and immune biology to human cancers. Here, we report the results from a dose-escalation clinical trial of intratumoral collagen-binding IL-2 and IL-12 cytokines in pet dogs with malignant melanoma, observing encouraging local and regional responses to therapy that may suggest human clinical benefit with this approach.

## MAIN

The recent success of immune checkpoint inhibitors has ushered in a new era to treat advanced cancers through rational engagement of the immune system^1–3^. Remarkable objective responses have been observed at primary tumors across a multitude of cancer immunotherapy strategies, although achievement of objective responses at metastatic sites remains an elusive clinical outcome for the majority of patients^4–6^. As such, combinations of checkpoint inhibitors with immune agonists have been explored to enhance systemic anti-tumor responses by overcoming immune-suppressive barriers operative at these metastatic sites^7,8^. In particular, the cytokines interleukin-2 (IL-2) and interleukin-12 (IL-12) have garnered significant interest owing to their ability to proliferate, activate, and differentiate critical effector immune cell populations unleashed by checkpoint inhibitors^9,10^. Encouraging synergy has been observed with these interleukin/checkpoint inhibitor combinations in early clinical trials, although adverse side effects have been encountered in patients^11–13^. As key signaling molecules between immune cells, endogenous immune-stimulating cytokines like IL-2 and IL-12 exhibit tightly controlled spatial distributions and diffusional kinetics to prevent aberrant and pathologic activation. However, in the therapeutic setting, systemically-dosed cytokines can elicit on-target, off-tumor activation of immune cells and subsequently possess an extremely narrow therapeutic window constrained by dose-limiting toxicities^14–16^. These clinical limitations resulting from cytokines administered systemically have driven recent interest in protein engineering strategies to mitigate systemic toxicities, through tumor-targeting immunocytokines^17–21^, conditionally-active/masked cytokines^22–26^, and receptor-biased cytokine agonists^27–30^ to enable their inclusivity alongside checkpoint inhibitors and other first-line cancer treatments such as radiation, chemotherapy, and surgery.

These elegant protein engineering efforts converge on the same objective for cytokine therapies: promote their accumulation within the tumor and constrain their signaling to the immediate tumor microenvironment. With advances in image-guided injection techniques, intratumoral dosing of therapies is now possible for the majority of solid tumor indications. As such, we and others have begun to explore strategies to physically retain cytokines like IL-2 and IL-12 within the tumor microenvironment after intratumoral injection through binding to co-dosed biomaterials^31–34^ or extracellular matrix components like collagen^25,35–37^. These approaches minimize the systemic biodistribution, tumor accumulation, and toxicity challenges associated with systemic dosing of engineered cytokines, and have led to marked improvements in both safety and efficacy profiles versus non-retained cytokines in mouse tumor models^31,35,36^. However, mouse syngeneic transplant tumor models lack the long-term immune selection pressures that sculpt human tumor genetics and thus they incompletely recapitulate critical evolutionary features of the complex human tumor microenvironment^38,39^. As a result, the achievement of treatment efficacy in mouse preclinical models with investigational immunotherapies is not sufficient for predicting their success when translated to human clinical trials^40–42^. For this reason, naturally-occurring tumors in larger companion animals complement these conventional model systems by illuminating the nuanced and complex tumor-immune interactions otherwise undetectable in mouse tumors, aiding translational investigation of novel anti-cancer strategies.

Here, we build upon our prior work in murine tumor models by examining the safety and efficacy of intratumorally-delivered, collagen-retained IL-2 and IL-12 cytokines in advanced malignant melanomas that spontaneously develop in outbred pet dogs. Dogs develop cancer at similar rates to humans, yet are an underutilized model to bridge the gaps between mouse and human studies of novel immunotherapies or treatment combinations^43–45^. Canine tumors feature many of the same biological immune escape mechanisms, driver mutations, and intratumor genetic heterogeneity that define human cancers, while also possessing more human-relevant body characteristics that enable prediction of drug biodistribution and PK/PD^46–49^. Moreover, a significant fraction of pet dogs with cancer presents with metastatic disease, enabling the evaluation of locoregional response to intratumoral therapy, which has been far more difficult to model and test in murine tumors or GEMMs. We previously evaluated the safety and mechanism of action of an intratumoral collagen-binding cytokine approach in canine soft tissue sarcomas, but did not have the opportunity to investigate long term anti-tumor responses due to the medical ethical obligation to resect such tumors shortly after treatment^50^. Guided by palliative regimens for malignant melanoma using hypofractionated radiation therapy (RT), we here report our studies of the safety and efficacy of a single RT dose with repeat dosing of tumor-localized IL-2 and IL-12 cytokines against malignant melanoma, a canine cancer that metastasizes in over 70% of cases^51^. Through a dose-escalation trial inclusive of key immunobiologic endpoints, we observed provocative activity engendered at both primary and metastatic tumors in a defined cohort of pet dogs. Profiling of canine patients that progress after therapy inform hypotheses regarding new therapeutic combinations predicted to improve tumor response rates, and we intend to deploy these strategies in both mouse models and pet dogs with naturally occurring cancers. Collectively, these efforts underscore the potential utility of comparative oncology inclusive of canine tumors to build, test, and optimize treatment regimens prior to commencing human clinical studies.

## RESULTS

### Patient enrollment and study population

For this study, clients whose dogs met trial inclusion criteria provided written informed consent before enrollment, and all procedures were performed in accordance with the study protocol approved by the University of Illinois Urbana-Champaign (UIUC) IACUC. Dogs were eligible after histologic or cytologic confirmation of oral malignant melanoma (OMM; n=14) or malignant melanoma involving other facial structures (n=1) and if their primary tumor was between 0.5-7.5 cm in diameter. Eligible dogs were also required to have adequate organ function as measured by standard laboratory tests, and have had a minimum three-week washout period if they had been recently treated with radiation therapy, systemic chemotherapy, immunotherapy, or any additional homeopathic/alternative therapy. There were no exclusion criteria for tumor stage or metastatic burden, age, weight, sex, breed, or neuter status for this study. Dogs were sequentially enrolled into a modified-Fibonacci 3+3 dose escalation trial design, with the initial IL-2 and IL-12 cytokine dose chosen from prior allometric scaling calculations and evaluation in both healthy beagles and pet dogs with soft tissue sarcomas (**Table 1**)^50^. In total, 15 dogs with median age 11 (min: 4, max: 16) were enrolled into the trial, with 10/15 (66%) dogs presenting with WHO Stage III or greater tumors, indicating metastatic disease at lymph nodes or lung tissue sites (**Extended Data Figure 1**).

**Table 1:** Dosing information and baseline patient characteristics. Description and dosing group allocation of 15 canine patients enrolled in the study. Patient breed, age, weight, tumor location, initial volume, and World Health Organization (WHO) domestic animal tumor stage are reported.

|  | Cohort 1x<br>(n=3) | Cohort 2x<br>(n=6) | Cohort 3.3x<br>(n=4) | Cohort 5x<br>(n=2) | Total<br>(n=15) |
| --- | --- | --- | --- | --- | --- |
| **Dosing Information** |  |  |  |  |  |
| LAIR-CSA-IL-2 dose ("IL-2"; µg/kg) | 17.4 | 34.8 | 57.4 | 87.0 | 80 doses |
| IL-12-CSA-LAIR dose ("IL-12"; µg/kg) | 2.08 | 4.16 | 6.86 | 10.4 | 80 doses |
| **Breed** |  |  |  |  |  |
| Purebred |  |  |  |  |  |
| Miniature Schnauzer | 1 (33%) | - | - | - | 1 (6.7%) |
| German Shepard | 1 (33%) | - | - | - | 1 (6.7%) |
| German Shorthaired Pointer | - | 1 (16.6%) | - | - | 1 (6.7%) |
| Labrador Retriever | - | 1 (16.6%) | 1 (25%) | 1 (50%) | 3 (20%) |
| Dachshund | - | - | 1 (25%) | - | 1 (6.7%) |
| Yorkshire Terrier | - | - | 1 (25%) | - | 1 (6.7%) |
| Shih Tzu | - | 1 (16.6%) | - | - | 1 (6.7%) |
| Standard Poodle | - | - | - | 1 (50%) | 1 (6.7%) |
| Australian Cattle Dog | - | - | 1 (25%) | - | 1 (6.7%) |
| Mixed Breed | 1 (33%) | 3 (50%) | - | - | 4 (26.7%) |
| **Primary site of malignant melanoma [n (%)]** |  |  |  |  |  |
| Lip/Buccal Mucosa | - | 2 (33%) | - | 1 (50%) | 3 (20%) |
| Mandible/Mandibular Mucosa | 1 (33%) | 3 (50%) | 1 (25%) | - | 5 (33.3%) |
| Maxilla/Maxillary Mucosa | 1 (33%) | 1 (17%) | 3 (75%) | 1 (50%) | 6 (40%) |
| Periocular | 1 (33%) | - | - | - | 1 (16.7%) |
| **Age (years)** |  |  |  |  |  |
| Median (min, max) | 11 (4, 13) | 11.5 (8, 16) | 10.5 (7, 12) | 10.5 (10, 11) | 11 (4, 16) |
| **Baseline weight (kg)** |  |  |  |  |  |
| Median (min, max) | 23.2 (5.8, 33.2) | 16.5 (6.8, 33.9) | 12.8 (4.7, 31.8) | 34.9 (29.3, 40.4) | 21.2 (4.7, 40.4) |
| **Baseline tumor volume (cm<sup>3</sup>)** |  |  |  |  |  |
| Median (min, max) | 7.5 (4.7, 11.6) | 6.8 (0.5, 16.3) | 7.9 (2.7, 18.6) | 23.2 (3.0, 43.4) | 7.5 (0.5, 43.4) |
| **Baseline WHO Stage [n (%)]** |  |  |  |  |  |
| I | - | 1 (17%) | - | - | 1 (6.7%) |
| II | 1 (33%) | 2 (33%) | 1 (25%) | - | 4 (26.7%) |
| III | 1 (33%) | 2 (33%) | 2 (50%) | - | 5 (33%) |
| IV | 1 (33%) | 1 (17%) | 1 (25%) | 2 (100%) | 5 (33%) |

### Tumor-localized IL-2/IL-12 with radiation is effective against canine oral melanoma

The primary objective of this study was to examine the anti-tumor efficacy potentiated by the combination of intratumoral collagen-anchored IL-2 and IL-12 with a single dose of radiation therapy. As current veterinary practice patterns favor the use of hypofractionated radiation therapy (RT) protocols using 8-10 Gray fraction size for OMM^52–54^, dogs treated in this study were provided a single RT dose of 9 Gy to stimulate tumor cell death and antigen generation. Local and regional lymph nodes were not irradiated, regardless of appearance or suspicion of possible metastatic disease. Dogs then received 6 doses of intratumoral collagen-anchored cytokines at the same two-week cadence similar to an existing FDA-approved intratumoral immune strategy (e.g. T-VEC) (**Figure 1a**). Pursuit of consecutive additional RT doses was not instituted due to concerns for detrimental lymphodepletion within the tumor and draining lymph node following preliminary experiments in the murine B16F10 model and other reports^55–57^ (**Extended Data Figure 2**). Moreover, the subsequent dosing of intratumoral cytokine alone enabled attribution of patient symptoms uniquely to cytokine treatment, and bypassed the requirement to deconvolute individual or interactive toxicities generated by continuous combinatorial therapy of RT with IL-2 and IL-12. All dogs were monitored for 48 hours after intratumoral cytokine dosing for symptoms of toxicity and had periodic blood draws performed for cellular and chemistry analyses.

**Figure 1.**
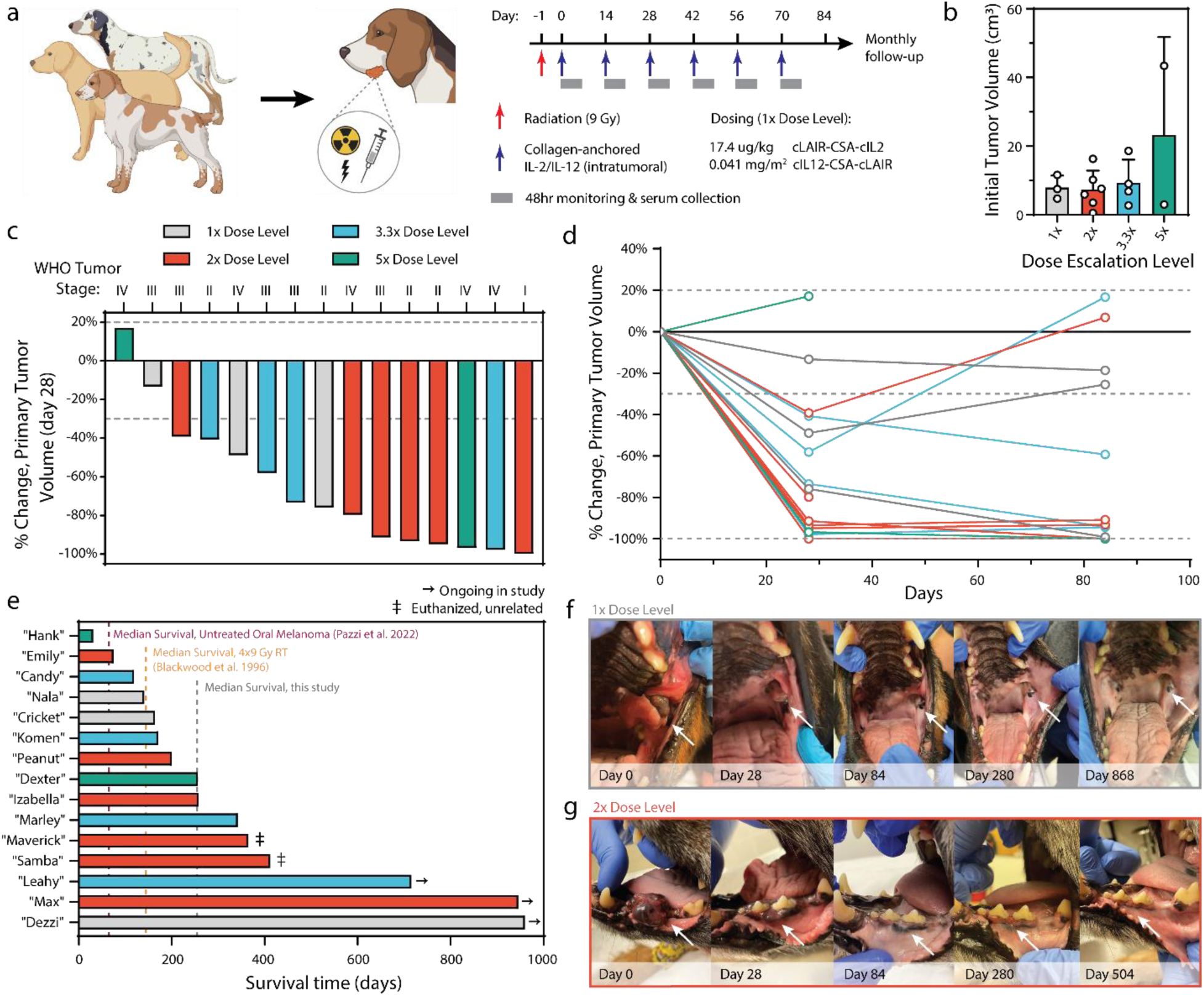
Study design and treatment outcomes. (a) Study-eligible dogs received 9 Gray (Gy) of radiation (red arrow) followed by 6 doses of intratumorally administered cytokines (blue arrows). Each cytokine dose was followed by 48 hours of clinical monitoring and serum collection. (b) Pretreatment primary tumor size quantified via CT radiologic assessment. (c) Percent change in tumor volume after radiation and 2 doses of intratumorally administered cytokines. Dotted lines depict RECIST criteria for tumor progression or clinical response. (d) Percent change in primary tumor volume over the course of treatment with intratumorally administered cytokines. One patient in each of the 2x and 5x dosing cohorts was euthanized prior to day 84 due to outgrowth of metastatic or primary tumors. (e) Swimmer plot of length of patient survival after trial start. (f-g) Images of primary tumors taken at indicated time points from select dogs from the 1x (f) and 2x (g) cohorts who displayed durable and complete response to treatment.

Primary tumor volumes at the time of first intratumoral dose had a median volume of 7.5 cm^3^ (min: 0.5, max: 43.4), although the highest dose cohort (‘5x’) included a dog with a primary tumor volume near the upper end of our eligibility criteria (**Figure 1b**). Responses to therapy were evaluated through comparative and serial assessments of computed tomography (CT) scans of primary tumor and associated regional metastatic lymph nodes identified at baseline (pre-treatment) with subsequent CT scans performed at day 28 and day 84. Rapid primary tumor volume reduction occurred in 13/15 (86.7%) malignant melanomas at the day 28 scan after just two doses of cytokine therapy and single RT dose (**Figure 1c**). At the day 84 CT scan performed two weeks after the final (6^th^) dose of intratumoral cytokine treatment, primary tumor responses were found to be stable or have further improved for 10/13 (76.9%) surviving dogs (**Figure 1d**). Two patients were euthanized before the day 84 CT tumor measurement due to progression of their primary and/or metastatic tumor sites. These tissues were collected for additional analysis detailed later in this study.

Treated pet dogs were followed after the twelve-week treatment period to monitor the durability of their responses and assess overall survival. As of the time of writing (January 2024), median survival regardless of tumor stage is 256 days, with three dogs still alive past two years (**Figure 1e, Extended Data Figure 3**). This is in contrast to reported median survival of 65 days for dogs with untreated oral melanoma^58^ and 147 days for OMM dogs treated with 9 Gy x 4 RT^53^. Two dogs were euthanized due to unrelated issues (age/quality of life; development of sinonasal chondrosarcoma) nearly a year after completing treatment. Interestingly, there appeared to be no correlation between the cytokine dose level and overall survival (**Extended Data Figure 4**). Of the dogs alive nearly 1000 days after treatment, the local response to therapy was rapid and robust, with less treatment morbidity than curative-intent surgical removal of OMM (**Figure 1f,g**). Overall, the objective responses observed in these canine patients with advanced stage and heterogeneous primary tumors were favorable, and further corroborate and extend upon the documented anticancer activities demonstrated in mouse models treated with the same collagen-binding cytokine approach^35,37^.

### Effective intratumoral doses of IL-2/IL-12 are also safe in pet dogs

The clinical promise of IL-2 and IL-12 cytokines has been limited by the toxicities observed at therapeutically effective doses^9,10,15,16,59,60^. As such, evaluating if the collagen-anchoring approach would ameliorate cytokine-driven toxicities at doses capable of promoting anti-tumor responses in pet dogs was paramount and translationally relevant. Analysis of whole blood at intervals following the first and second doses of intratumoral cytokine therapy indicated minimal elevation of systemic alanine transaminase (ALT) levels for most patients tested at the lowest three dose levels, with ALT levels normalizing prior to administration of each subsequent cytokine dose (**Figure 2a**). The predominant adverse events observed were mostly grade 1 and 2 across dose-level cohorts, with the most commonly occurring events being associated with hemoglobinemia, thrombocytopenia, lethargy, anorexia, and elevation in ALT and ALP levels (**Extended Data Figure 5**). The owner of one 2x-dose-level dog with elevated ALT chose not to pursue the 6^th^ dose of cytokine treatment. Select dogs demonstrated elevated ALT in the 3.3x dose cohort and responded well to s-adenosylmethionine and silybin to mitigate hepatocyte toxicity and normalize liver function. More clinically significant ALT elevation and symptoms consistent with cytokine release syndrome (i.e. thrombocytopenia, hypoproteinemia, severe lethargy, pyrexia) were observed in the 5x dose cohort (**Extended Data Figure 5**). These patients received supportive care including intravenous fluids and dexamethasone SP (0.5 mg/kg, IV) and fully recovered after treatment. A reduction in subsequent doses to this cohort to 3.3x was instituted to minimize discomfort and health risks in these patients.

**Figure 2.**
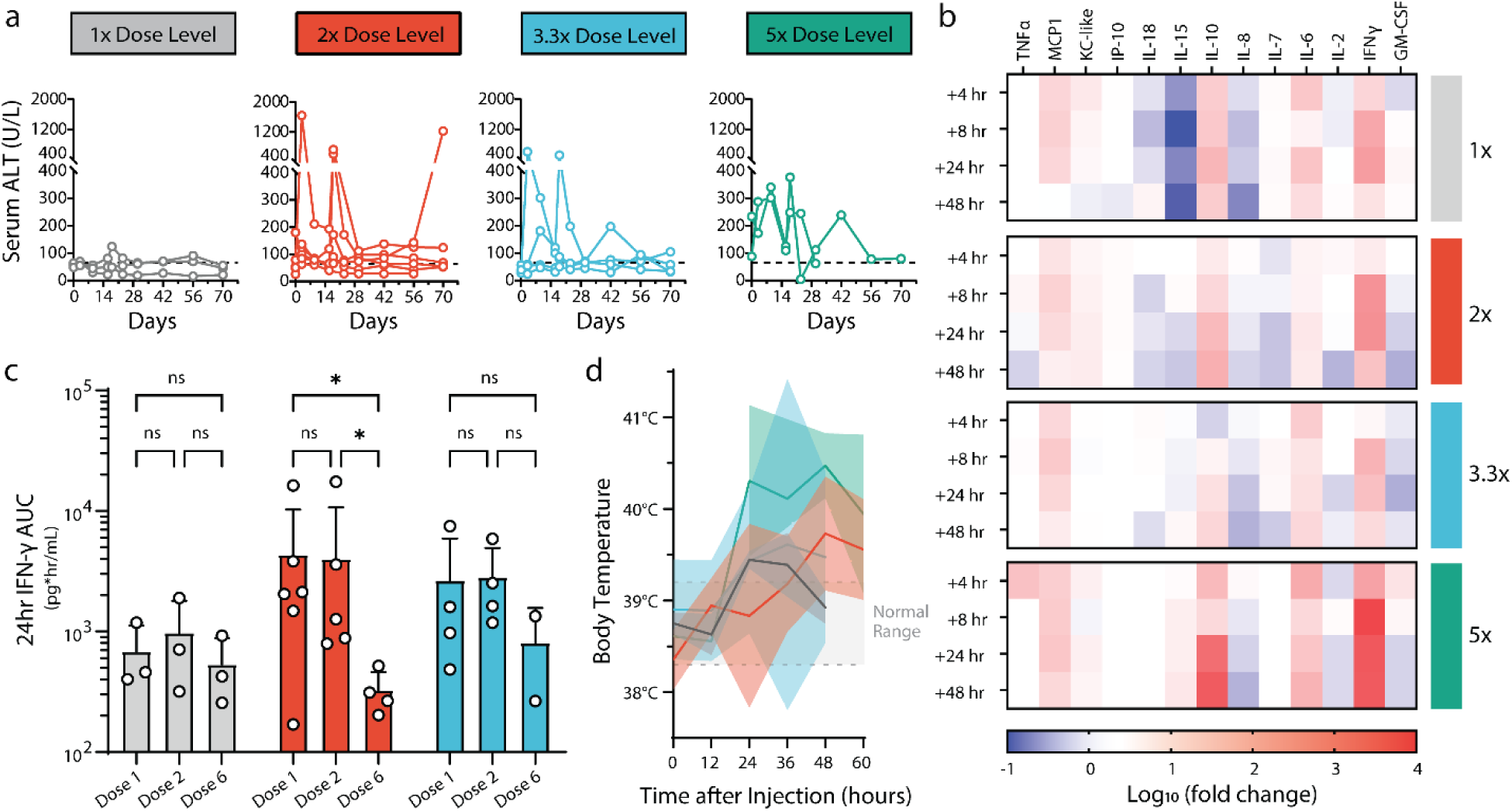
Safety profile of collagen-anchored cytokine therapy. (a) Serum alanine transaminase (ALT) levels measured via blood work at indicated time points following intratumoral cytokine dosing at day 0 and day 14. Dotted line indicates a clinically healthy ALT threshold. (b) Serum was collected at several time points after the first intratumoral cytokine injection and analyzed for cytokines and chemokines. Heatmap rows describe averaged sera data from each dosing cohort, reported as log_10_ fold change in concentration compared with pretreatment values. (c) Serum was collected 4-, 8-, and 24-hours post-cytokine administration after the indicated doses and analyzed for systemic exposure to interferon gamma (IFN-γ), as represented by 24-hour IFN-γ area under the curve (AUC). (d) Body temperature of patients was measured at the indicated time points after the first cytokine administration. Dotted lines indicate normal body temperature range. Statistics: IFN-γ AUCs compared by two-way ANOVA with Tukey’s multiple comparisons test. ns, not significant; *P < 0.05.

To correlate the observed clinical activity and potential toxicity with pharmacodynamic biomarkers, profiling of the systemic chemokine/cytokine responses to combination RT with intratumoral cytokine treatment was performed. Similar response dynamics to those previously reported were observed, wherein IL-12 drives elevation of systemic levels of interferon gamma (IFN-γ), with a delay in the elevation of IL-10 (**Figure 2b**) ^50,61–64^. Peak levels of IFN-γ were mostly consistent among the lowest three dose cohorts, but spiked significantly higher at the more toxic 5x dose level. To confirm circulating elevations of IFN-γ and IL-10 were biomarkers of intratumoral cytokine activities and not an epiphenomenon of ionizing radiation or injection site trauma, an additional cohort of four dogs receiving only a single dose of RT (9 Gy) and sham intratumoral saline injection was analyzed, and no measurable concentrations of circulating IFN-γ or IL-10 was identified (**Extended Data Figure 6**). Moreover, a cohort of three dogs receiving intratumoral cytokine only without RT demonstrated similar dynamics of IFN-γ and IL-10 changes following treatment, providing further evidence that the dynamic responses observed via multiplex-serum profiling are IL-2/IL-12 mediated rather than due to the combination of RT with intratumoral cytokine treatment (**Extended Data Figure 6**).

Given the importance of IFN-γ both directly on tumor cells and in facilitating productive anti-tumor immune responses^65–68^, an estimation of the systemic exposure of patients to IFN-γ via area-under-the-curve (AUC) was calculated. The analysis provided some evidence of immune tachyphylaxis, in which the response to intratumoral cytokine therapy appears to have diminished by the sixth dose, relative to the responses to the initial doses of therapy (**Figure 2c**) This is most pronounced in the 2x dose cohort, although some pet owners elected to not continue treatment with 6 doses of intratumoral cytokine therapy due to complete regression of the local tumor site concurrent with some adverse toxicities (2 of 4 dogs, 50%), confounding the statistical comparisons at the 3.3x dose level. The phenomenon of immunologic defervescence has been difficult to study in murine models, but has been noted in human patients, highlighting the potential utility to examine various treatment regimens in dogs to minimize tachyphylaxis. Characterization of anti-drug antibody responses that could attenuate immunostimulatory activities to collagen-anchored cytokines found the existence of antibodies but not at levels high enough to explain the magnitude of reduced IFN-γ response at the final dose timepoint (**Extended Data Figure 7**).

Finally, patient body temperatures were measured during the post-treatment monitoring phase, and it was observed that most dogs became mildly febrile regardless of dose level (**Figure 2d**). These mild symptoms did not require medical intervention, and were often accompanied by transient inappetence and lethargy amongst patients during the monitoring phase. Overall, the responses potentiated by therapy were well-tolerated at the 1x and 2x dose levels, with dose-limiting toxicities first observed at the 3.3x dose level in a subset of patients but amongst a majority of patients at 5x.

### Tumor-localized IL-2/IL-12 with RT potentiates responses at metastatic lesions

Many pet dogs enrolled in this trial presented with metastatic lesions, providing an opportunity to examine whether local treatment of the primary tumor with IL-2, IL-12, and RT could promote locoregional responses at untreated metastatic sites, an important outcome for intratumoral therapies. CT measurements were obtained for metastatic lymph nodes and measured for radiologic response in comparison with their pre-treatment volumes (**Figure 3a**). Following treatment of primary tumors with RT and intratumoral cytokines, 3/10 dogs (30%) displayed a partial response at metastatic lymph nodes. Two additional dogs achieved stable disease during the treatment period, for an overall biologic response rate to combination therapy of 50% (5/10 dogs). Two dogs were euthanized prior to the day 84 measurement; one due to suspected progression of brain/CNS metastases, and another for significant progression of lung metastases. For a subset of the responding patients, appreciable regional edema was present at metastatic lymph node sites at the interim (day 28) measurement.

**Figure 3.**
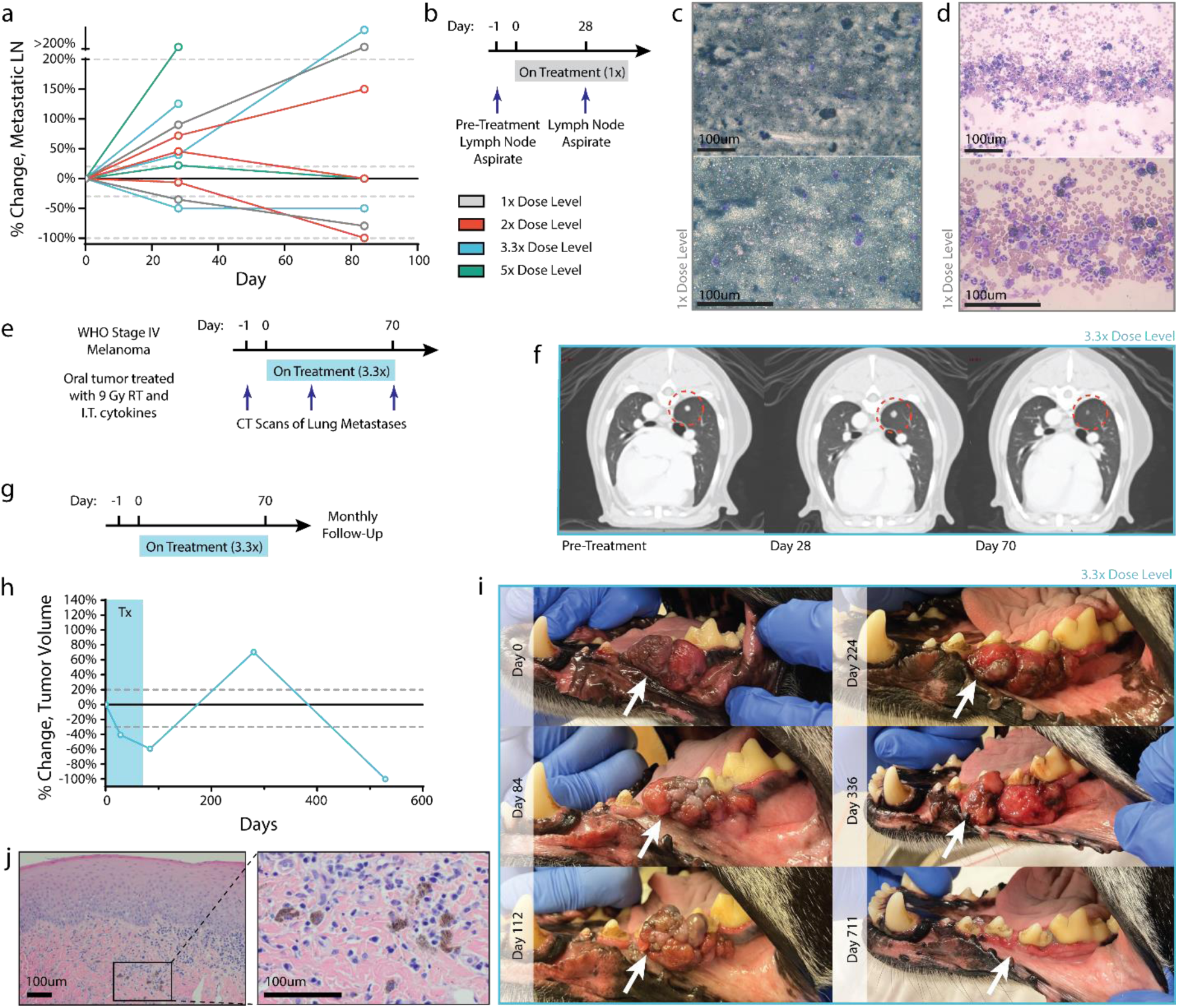
Case studies of patients demonstrating abscopal immune responses. (a) Percent change in volume of regional lymph node metastasis relative to pre-treatment volume, as determined by CT measurement. (b) Fine needle aspirates were collected from the lymph node of a patient in the 1x cohort before treatment and after 2 intratumoral cytokine doses. (c) Pretreatment aspirate shows diffuse infiltration of melanocytes. (d) Lymph node disease is decreased after 2 cytokine treatments, with a marked increase in polymorphonuclear immune cells. (e) CT images from a stage IV patient in the 3.3x treatment group were collected tracking the progression of a lung metastasis after local treatment of oral melanoma. (f) CT images suggest pseudoprogression of a lung metastasis after early cytokine doses, with later regression after additional cytokine doses. (g-i) A patient in the 3.3x dosing group received a full course of treatment and had routine follow-up visits to monitor tumor progression. Tumor measurements (h) and images (i) were taken at the indicated time points, demonstrating a significantly delayed treatment response. (j) Hematoxylin and eosin staining on this tumor showed an absence of tumor cells with only scattered melanophages observed at day 529. Scale bars: 100um.

For one patient, a pre-treatment fine needle aspirate (FNA) of the tumor-draining lymph node as well as a subsequent FNA of the same regional lymph node on the day 28 CT scan were obtained (**Figure 3b**). Prior to treatment, this lymph node was completely effaced with disease, as detected via the absence of immune cells and the majority presence of cancerous melanocytes and extracellular melanin (**Figure 3c**). After two doses of intratumoral cytokine and single RT treatment, the lymph node CT scan indicated a robust decrease in metastatic regional lymph node volume (−35.3%; partial response) and concurrent immunologic clearing of melanoma cells and pigmentation (**Figure 3d**). We observed the presence of polymorphonuclear (PMN) cells, likely neutrophils, in the FNA, many of which had phagocytosed tumor cell debris and melanin. One additional patient had detectable lung metastasis at time of presentation and trial enrollment (**Figure 3e**). While this dog ultimately succumbed to progressive metastatic disease, there was evidence of at least one regressing lung metastasis lesion during treatment (**Figure 3f**). This mixed abscopal response may be due to underlying genetic differences between primary and disseminated disease, as well as among differing clonally-derived lung metastases^69–71^. However, the locoregional response of metastatic disease to combined intratumoral IL-2/IL-12 and single-dose RT treatment is consistent with an immune-mediated mechanism of action, and similar to prior reports of combined radiation with immunotherapy^72–76^.

Similar to the pivotal Phase III clinical trials with T-VEC^77^, out of concern that longitudinal sampling of the primary treated tumors could confound results by introducing additional paths for intratumoral dose egress, we did not profile the immune response to therapy during treatment. However, building upon our prior characterization of the immune-mediated response to collagen-anchored cytokines in canine soft tissue sarcoma and murine tumors^35,50^, we highlight an anecdotal case of long-term anti-tumor response after the completion of treatment in oral melanoma which presumably involved immune activity.

One patient had a strong primary tumor response while on-therapy, but displayed slow growth of that tumor in the year following treatment completion (**Figure 3g**). However, at the 12-month follow-up appointment after treatment, the primary tumor was no longer visible and was later confirmed to be absent via CT (**Figure 3h-i**), as well as histopathology (**Figure 3j**).

Additional immunohistochemistry for Melan-A further confirmed the absence of disease in the gingival tissue of this patient at day 529 (**Extended Data Figure 8**). While examples of spontaneous human tumor regressions have been reported^78,79^, they are quite rare (∼10^-5^)^78^. The slow post-treatment tumor growth may correspond to a state of immune equilibrium, leading eventually to tumor elimination, similar to other immunotherapy approaches^80,81^.

### Dysfunctional antigen presentation predicts resistance to tumor-localized cytokine therapy

Identifying and understanding which factors, if any, contributed to poor response to the combined RT plus intratumoral cytokine treatment regimen was further studied. Towards this goal, FFPE-processed primary and metastatic tumor tissue from eight dogs who were euthanized for progressive disease were advanced for detailed histologic and genomic evaluations. No clear trends were observed between overall survival of these progressor patients and immune infiltration status profiled through immunohistochemistry for CD3 and Iba1 (**Extended Data Figure 9**). Extracted RNA from these tissue sections were profiled using the Nanostring nCounter platform (**Figure 4a**). A hierarchical cluster of pathway-specific gene expression emerged that encompassed the coordination of innate and adaptive immunity, including T-cell, B-cell, and macrophage function as well as antigen presentation (**Figure 4b**). Within this cluster, varied expression amongst the progressor dogs was observed, and additional unsupervised clustering of the antigen presentation gene set yielded two clusters of 4 dogs each (**Figure 4c**). Given that tumor dysregulation of antigen presentation and response to IFN-γ is a common immune evasion mechanism^66,82,83^, the expression of MHC class I-related genes were examined, and identified a significant difference in *B2m* and *Dla-79* transcripts between the clusters of progressor dogs (**Figure 4d**). This result suggested that the first cluster of dogs may have had impaired MHC-I expression, at least amongst a partial population within the heterogeneous tumor. Broader comparisons in gene expression between these two cohorts indicated greater expression of effector lymphocyte-associated genes such as *Slamf6*, *Ctsw*, and *Trgc3* as well as interferon-inducible genes including *Ido1*, *Gbp5*, and *Cxcl10* amongst the MHC-I higher expression cohort, Cluster 2 (**Figure 4e**).

**Figure 4.**
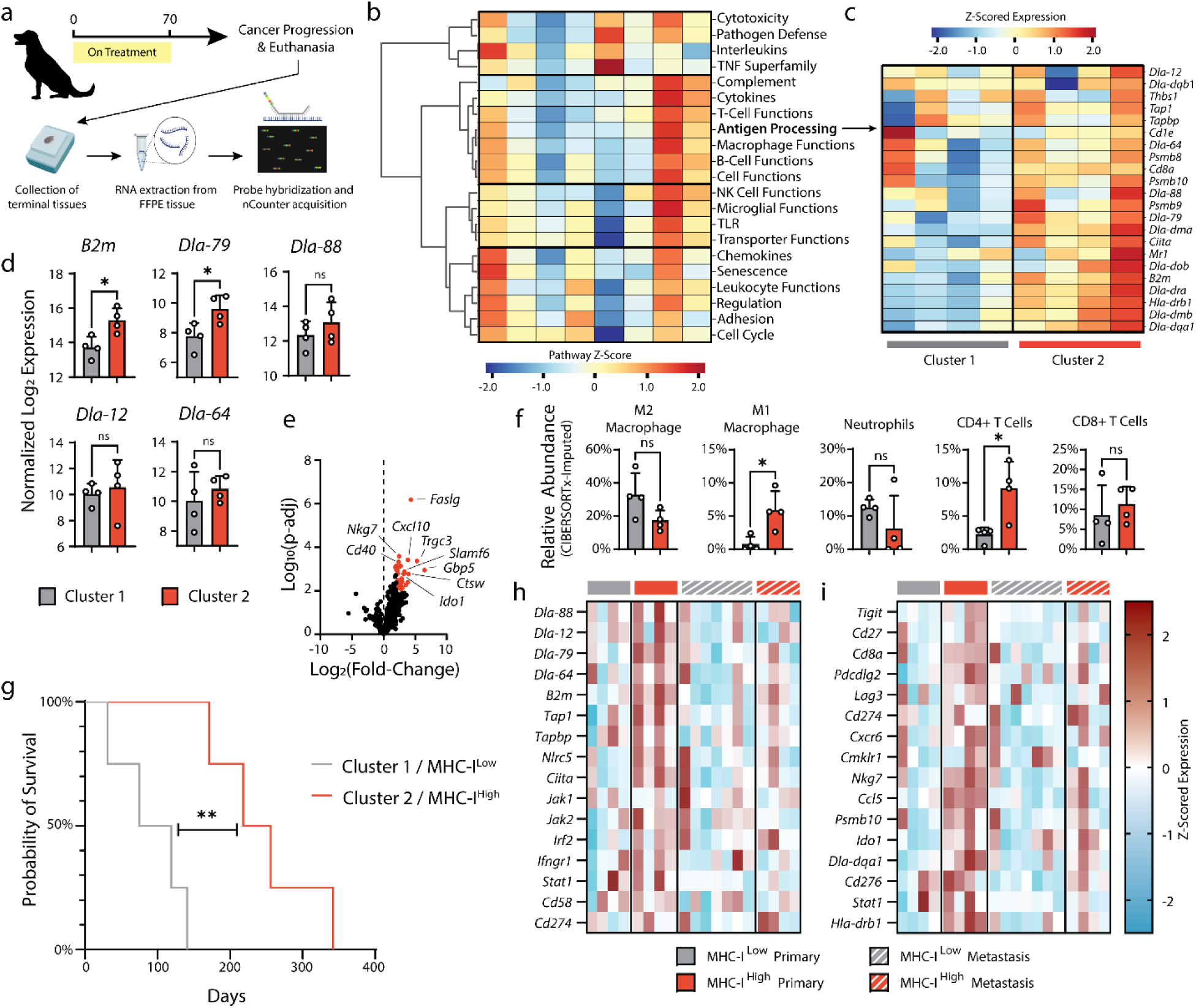
Nanostring RNA profiling of terminal primary and metastatic tumor tissues. (a) Terminal primary and metastatic tumor tissues from euthanized patients were collected and FFPE processed. RNA was extracted from FFPE tissues and prepared for NanoString analysis with the NanoString Canine ImmunoOncology nCounter panel. (b) Pathway scoring and hierarchical clustering of NanoString annotated pathways involved in canine cancer immune response. Pathway scores were calculated as the first principal component of the pathway genes normalized expression. Heatmap columns represent individual patients’ primary oral melanoma. (c) Z-scored expression of genes related to canine antigen presentation, with tumor samples grouping into two hierarchical clusters. (d) Normalized expression (log 2) of MHC class-I related genes. (e) Volcano plot of differential gene expression of cluster 2 (MHC-I^Hi^) relative to cluster 1 (MHC-I^Low^). Genes associated with significant P-adj values (<0.05) are highlighted in red. (f) Relative abundance of intratumoral immune populations as determined through application of the CIBERSORTx algorithm on NanoString data. (g) Survival of MHC-I^Hi^ and MHC^Low^ progressor dogs. (h-i) Z-scored expression data for genes associated with tumor immune escape^71^ (h) and response to immune checkpoint blockade^86^ (i) for primary and metastatic lesions of MHC-I^Hi^ and MHC^Low^ patients. Statistics: Differential gene expression and relative abundance of immune populations compared using one-way ANOVA with Tukey’s multiple comparisons test. Survival compared with log-rank Mantel-Cox test. ns, not significant; *P<0.05; **P<0.01.

Intriguingly, the most differentially expressed gene was for Fas-ligand (*Faslg*) and may represent a consistent mechanism of immune escape within the cohort of progressor dogs (Cluster 2) with greater *B2m* expression. It has been established that peripheral expression of Fas-ligand on multiple cell types in response to inflammatory stimulus promotes deletion of auto-reactive T lymphocytes (e.g. peripheral tolerance)^84^, so we examined whether there were compositional differences in the immune compartments from the tumors of the progressor dog cohorts. Using CIBERSORTx^85^, the relative abundance of immune cell populations from the bulk Nanostring profiling data were estimated. Tumors with reduced *B2m* expression were accompanied by greater populations of canonical tumor-suppressive immune cells (i.e. “M2” polarized macrophages, neutrophils), while dogs with higher MHC-I antigen presentation had more activated macrophages and CD4 T lymphocytes (**Figure 4f**). Together, these differences likely contributed to the poorer prognosis of patients with reduced MHC class I antigen presentation, regardless of tumor stage at presentation (**Figure 4g**, Log-rank hazard ratio: 4.472).

To explore why the cohort of dogs with higher class I antigen presentation and reduced abundance of immunosuppressive immune populations (Cluster 2) still progressed after therapy, gene expression was examined within tissue collected from metastatic tumor sites. Using a gene set describing common genetic mutations that enable immune escape at primary or metastatic tumor tissues^71^, the differences in expression between cohorts is diminished at the metastatic tumors (**Figure 4h**). This suggests that the metastatic tumors from dogs with higher MHC-I expression at their primary tumors may have been preferentially seeded by tumor subpopulations with greater genetic immune escape, such as MHC-I loss of heterozygosity. We further examined gene expression between cohorts at their primary and metastatic tumors across an annotated set predictive of human response to checkpoint inhibitors^86^. We found that only the primary tumors of higher expression MHC-I dogs are expected to have positive response to immunotherapy, consistent with the observed local response but metastatic progression of these patients following our combined cytokine treatment (**Figure 4i**). Overall, these results motivate exploration of treatment combinations to overcome dysfunctional MHC class I antigen presentation in tumors to extend therapeutic benefit to a greater population of pet dogs, with the intention that lessons gleaned from comparative oncology studies can be quickly pivoted to accelerate novel immunotherapeutic strategies to benefit human cancer patients.

## DISCUSSION

Mechanisms of primary and adaptive resistance to immunotherapy contribute to the lack of clinical benefit for a majority of cancer patients treated with antagonistic, checkpoint-inhibiting antibodies^5^. As a result, there have been attempts to combine these therapies with agonistic, or immune-stimulating, agents to overcome tumor resistance mechanisms and drive more durable responses^87,88^. Cytokines, such as IL-2 and IL-12, are one class of agonistic therapies that have shown great promise against human cancers, but suffer from unacceptable toxicities due to their activation of immune cells throughout the body^15,16^. Approaches to restrict the activity of potent cytokines to the tumor have gained momentum, one of which includes the retention of engineered cytokines to tumor extracellular matrix following intratumoral injection^25,35–37^. We and others have previously reported on the safety improvements provided by this strategy of anchoring cytokines to tumor collagen in both mice and pet dogs^25,35,50^, but the efficacy in advanced canine tumors was previously unexplored.

In this work, we evaluated the efficacy of tumor-localized IL-2 and IL-12 cytokines in pet dogs with advanced oral malignant melanoma to potentially predict success of clinically translating this approach. As dogs share key physical features and tumor biology with humans, they have gained traction as models for human comparative oncology^43,45,48^. Here, we have observed encouraging results for both the anti-tumor efficacy and tolerability of single-dose radiation therapy with repeat intratumoral IL-2 and IL-12 cytokines. Primary tumor responses were often rapid and durable, with 256-day median survival across all treated cohorts; significantly longer than the historical 65-day median for untreated canine oral melanoma^58^. Moreover, many of these responses were observed among dogs in the non-toxic 1x and 2x cohorts, suggesting that the tumor-localization strategy via retention to tumor collagen is clinically promising for safely and effectively treating human malignancies. Locoregional responses at metastatic sites driven by intratumoral therapy achieved an overall biologic response against combined tumor and metastases in 10/13 dogs (76.9%) receiving the full therapy, with partial responses in 8/13 (61.5%) of dogs (**Extended Data Figure 10**). This result provides early evidence that intratumoral treatment with collagen-bound cytokines may potentiate systemic anti-tumor immunity in pet dogs with naturally occurring cancers. Importantly, these canine tumors develop under evolving tumor immune evasion and suppression mechanisms analogous to those in humans, suggesting this engineered cytokine approach may achieve similar responses in human clinical trials.

Profiling of dogs that progressed while, or soon after, receiving the RT plus intratumoral cytokine treatment revealed that dysfunctional antigen presentation may contribute to the rapid progression of canine malignant melanoma. This complements a growing list of canine tumor features that overlap with the human hallmarks of cancer, including sustained proliferative signaling^89^, and mutations to oncogenic driver or tumor suppressor genes^49,90^. While less definitive, the dogs with higher MHC class-I expression may have progressed due to tumor microenvironment-induced dysfunction of immune cells. With the observation that *Faslg* and *Ido1* are more highly expressed by these tumors, we suspect that the combination cytokine therapy was actively promoting an anti-tumor response met by immune counter-regulation, as we observed previously in canine soft tissue sarcomas^50^. The combination of IL-12/IL-2 has been described to upregulate the expression of Fas-ligand on draining lymph node lymphocytes^91^, which, while aiding their ability to kill malignant tumor cells, could contribute to eventual lymphocyte fratricide or suicide^92^. This mechanism might contribute to our observation of tachyphylaxis in some of the dogs (**Figure 2c**). Moreover, the mixed response between primary tumors and metastatic sites may manifest from the varied gene expression landscape and erected barriers to immune function observed between these metastatic tumors and their primary tumor counterparts, suggesting that systemic therapies (such as anti-PD-1 antibodies) may be necessary to leverage cytotoxic effector cells primed by local intratumoral therapy^35,50^.

Our learnings from each group of progressing dogs provides actionable insights for future combination treatments to test alongside the intratumoral cytokine approach. To this end, we are interested in evaluating the combination of checkpoint inhibitors with the RT plus intratumoral cytokine treatment in future studies. Our prior work with canine soft tissue sarcomas indicated that checkpoint blockade might relieve counter-regulatory responses to intratumoral cytokine therapy, which we confirmed in the murine B16F10 tumor model^50^. However, resistance to intratumoral IL-2 and IL-12 therapy via beta-2-microglobulin (B2M) loss and subsequently, dysfunctional antigen presentation, appears to overlap with known resistance mechanisms to checkpoint inhibitors^82,93^. As a result, future screening of canine *B2m* and MHC class-I associated genes expression prior to trial enrollment could help accrue patients into separate, more rationally-designed combination treatments. For dogs with reduced or dysfunctional antigen presentation, there have been strategies reported for combining immunotherapies with epigenetic drugs to remove silencing of B2M and restore MHC-I expression^94–96^, in addition to strategies to engage innate immune cells for direct tumor-cell killing^97–99^ or to coordinate their licensing of antigen-independent killing by CD8+ lymphocytes^100,101^. Finally, given our observation of tachyphylaxis in response to repeat cytokine dosing and reports of the importance for immune rest in engineered CAR-T therapies^102^, we are interested in exploring longer intervals between cytokine doses to minimize AICD or induced dysfunction of primed CD8+ T cells.

Overall, this work highlights the benefit of pre-clinical evaluation of a novel immunotherapy alongside current standard of care in a more human-analogous cancer model than mouse tumors. While statistical power of such a trial in pet dogs is more limited, we argue that the value gained in predictive efficacy, safety, and resistance to therapy are obtained at dramatically lesser expense and greater speed than a corresponding human clinical trial. Exploitation of canine trials as a bridge from murine studies to the clinic should be expanded to reap these benefits more widely. Certain methodology to maximize value from these canine cancer trials stands to gain from broader investigation as well. We recognize that a primary limitation of this study is the lack of longitudinal sampling from canine tumors to characterize the evolution of anti-tumor responses as well as resistance to cytokine treatment. Through comparative oncologic testing, we anticipate a greater likelihood of future clinical success for our collagen-binding cytokine approach, as well as more broadly for other novel immunotherapies investigated in pet dogs with cancer.

## METHODS

### Ethics statement

This study complies with all relevant ethical norms and principles. This research study protocol was approved by the Institutional Animal Care and Use Committee at the University of Illinois Urbana-Champaign.

### Trial eligibility and enrollment of pet dogs

Client-owned pet dogs with cytologically or histologically confirmed OMM were included in the study. Eligibility criteria required dogs to have 1) primary tumor measure between 0.5 to 7.5 centimeters in diameter, 2) adequate organ function determined by laboratory evaluations (complete blood count, serum biochemical profile, and urinalysis), and 3) a minimum three-week washout period for radiation therapy, systemic chemotherapy, or any additional immunosuppressive/homeopathic/alternative therapy. No exclusion criteria for tumor stage or metastatic burden, age, weight, sex, or neuter status were applied for this trial. Tumor staging at enrollment was determined based the World Health Organization (WHO) staging scheme for dogs with oral melanoma^103^. All patient owners provided written consent before enrollment and all procedures were performed in accordance with the study protocol approved by the University of Illinois Urbana-Champaign (UIUC) IACUC.

### Collagen-anchoring IL-2 and IL-12 cytokine protein production

Canine cytokines (cLAIR-CSA-cIL-2, cIL-12-CSA-cLAIR) were cloned and recombinantly expressed as previously described^50^. Briefly, stable HEK293-F cell lines for each cytokine were prepared through cloning into the expression cassette of PiggyBac (System Biosciences) transposon vector, followed by dual transfection of the transposon vector and the Super PiggyBac transposase plasmid. Stable integration was confirmed after sorting EGFP+ cells 3-4 days after transfection (BD FACS Aria). Protein was produced from IL-2 and IL-12 expressing stable lines during one-week culture in serum-free media (Freestyle 293, Invitrogen) and purified with HisPur Ni-NTA affinity resin (ThermoFisher Scientific). Protein was analyzed by size exclusion chromatography (Superdex 200 Increase 10/300 GL column, Cytiva Life Sciences on AKTA FPLC system) for size and aggregation and validated to meet low endotoxin levels (<5EU/kg) by Endosafe Nexgen-PTS system (Charles River Labs). Activity of cytokines was confirmed through CTLL-2 and HEK Blue IL-12 activation assays, while collagen-binding was confirmed through ELISA. Aliquots of cytokines were snap-frozen in liquid nitrogen and thawed immediately prior to dilution in sterile saline for dosing intratumorally to dogs.

### Study design and intratumoral dosing of cytokines

Fifteen eligible dogs were enrolled into a modified-Fibonacci 3+3 dose escalation trial design of four different cohorts. The trial consisted of a regimen involving treatment with a single 9 Gray (Gy) dose of radiation therapy followed by 6 doses of cLAIR-CSA-cIL2 (IL-2) and cLAIR-CSA- cIL12 (IL-12) every two-weeks (Table 1). Radiation was delivered using a Varian™ TrueBeam™ linear accelerator with 6 MV photons at standard dose rate of 6 Gy/minute (Varian Medical Systems, Palo Alto, CA, USA). Depending on location and proximate organs at risk, dose was delivered either using manual calculations for parallel opposed portals, or with 3-dimensional conformal radiation plan using CT guidance and a treatment planning system (Varian Eclipse v.15). The dose was calculated to the central axis for parallel opposed portals, and with the goal of 100% of dose to 95% of the planning target volume (gross tumor volume plus a 3-5 mm expansion) for computer plans. The initial doses of IL-2 and IL-12 cytokines were determined from prior allometric scaling calculations and evaluation in both healthy beagles and pet dogs with soft tissue sarcomas^50^. Doses of cLAIR-CSA-cIL2 (17.4 μg/kg) and cIL12-CSA-cLAIR (2.08 µg/kg) were prepared from frozen protein aliquots and combined in a total volume not exceeding 0.5 mL in sterile saline. A 29-gauge, ½-inch insulin syringe was used to slowly inject the full dose volume via a single insertion point using a fanning pattern into the tumor. No additional measures were used to avoid any internal necrotic areas within the tumor. Radiation therapy was performed using Varian TrueBeam system. Adverse events were classified and graded in accordance with the Veterinary Cooperative Oncology Group’s Common Terminology Criteria for Adverse Events (VCOG-CTCAE v2)^104^.

### Clinical response assessment

Clinical and vital evaluations were conducted on all patients at baseline and preceding each treatment administration at the UIUC Veterinary Teaching Hospital. In addition, after intratumoral cytokine administration, a 48-hour monitoring period was initiated to assess the presence of any toxicity-related symptoms, coupled with blood sampling for complete blood count, serum biochemical profiling, and urinalysis. In addition, after each treatment, blood draws by jugular venipuncture were performed for cytokine/chemokine analysis before treatment, 2, 4, 8, 24, and 48 hours post treatment. Patients were followed-up until death or removal from the trial.

Clinical and caliper measurements of the maximum tumor and lymph node dimensions were conducted by board-certified veterinary oncologists and measurements were documented in millimeters during each examination. In addition, primary tumor or metastatic lesions were assessed by computed tomography (CT) (Somatom Definition AS, Siemens) at pre-treatment, day 28, day 70, and day 84 (two weeks after the last treatment). The tumor size and the percentage of change were determined based on CT measurements. Because determination of longest dimension is challenging with these frequently irregularly marginated tumors, tumor volume was used to measure response to therapy. Volume was determined using radiation therapy treatment planning software (Eclipse v15, Varian, Palo Alto, CA) by importing CT scan images (1.5 mm slices) before and after treatment. Gross tumor volume was delineated based on distortion of normal tissues by the mass effect combined with changes in Hounsfield units that reflect contrast enhancement due to changes in electron density. The software will yield a three dimensional volume based on the contours that are created. Standard criteria for volumetric assessment of tumor response were used. Furthermore, the assessment of tumor response was carried out during each visit and was determined in accordance with the guidelines established by the Response Evaluation Criteria for Solid Tumours in Dogs (v1.0) (VCOG)^105^. Patients presenting with stable or progressive disease were allowed to remain in the study under the condition that no adverse events were observed, or if such events could be mitigated through the implementation of a dose reduction protocol. Clients had the option to remove their dogs from study if their pets’ conditions worsened, they showed signs of declining health, or if the treatment caused unbearable side effects. The decision could be made by the investigator, the dog’s owner, or both.

### Multiplex cytokine assay and ELISA

Serum samples collected from patients following treatment were examined for concentrations of 13 cytokine and chemokine analytes, including GM-CSF, IFN-γ, IL-2, IL-6, IL-7, IL-8/CXCL8, IL-10, IL-15, IL-18, IP-10/ CXCL10, KC-like, MCP-1/CCL2, and TNFα (Canine MILLIPLEX Magnetic Bead Panel, Millipore Sigma) at Eve Technologies (Calgary, AB, Canada). Individual analyte concentrations were determined from panel standard curves for each cytokine or chemokine. Time course analysis of patient response to IL-2/IL-12 and radiation therapy was performed by determining the log10 fold-change of analyte concentrations relative to their pre-treatment levels. IFN-γ and IL-10 serum concentrations following treatment with collagen-anchored cytokines and radiation therapy were further measured using the Canine IFN-γ Quantikine ELISA kit (R&D) and the Canine IL-10 Quantikine ELISA kit (R&D) according to the manufacturer’s instructions.

### Nanostring RNA profiling

RNA was isolated from 10-μm FFPE samples from resected canine primary melanoma tumor or metastatic tumor lesions using an RNEasy FFPE Kit and deparaffinization solution (Qiagen). Isolated RNA was examined by Bioanalyzer (Agilent) for assessment of fragment size prior to hybridization with nCounter probe sets (Nanostring). Canine RNA samples were hybridized with the Canine IO nCounter Panel code set for 22 hours at 65°C per the manufacturer’s instructions. Following hybridization, samples were loaded into the analysis cartridge and scanned at maximum resolution using NanoString PrepStation and Digital Analyzer.

Canine RCC count files were normalized using nSolver software (Nanostring) after background thresholding using the mean of 8 negative control probes and batch correction against a panel standard control. Normalized gene counts were processed using the nSolver Advanced Analysis module for differential expression and pathway enrichment analysis. *P* value adjustment was performed using the Benjamini–Hochberg method to estimate FDRs of differentially expressed genes (DEG).

### Estimation of tumor immune cell abundance

Relative abundance of tumor-infiltrating cell fraction was estimated from bulk NanoString profiling data by employing CIBERSORTx^85^ algorithm using a validated leukocyte gene signature matrix (LM22). Bulk NanoString profiling data was assessed in relative mode, with 100 permutation runs and without quantile normalization.

### Immunohistochemistry and cytology

Canine advanced malignant melanoma tumors were resected at specific indicated timepoints. Following resection, tumor tissues were fixed in 10% formalin and subjected to a paraffin processing and embedding protocol. Immunohistochemistry (IHC) was used to determine the presence of inflammatory cells, specifically positive for CD3 (T lymphocyte; Biocare CP215C), Iba-1 (macrophage; Biocare, catalog no. CP 290 B, RRID:AB_10583150), and Melan-A (melanoma-specific antigen; Biocare A103) for melanoma cells. All samples were histologically evaluated and classified by a single board-certified veterinary pathologist. Tumor tissues were classified based on CD3 T cell infiltration status into an immune phenotype, defined as a) inflamed - highly infiltrated by CD3+ T cells, b) immune desert/cold – devoid of CD3+ T cells, and c) immune excluded – bordered yet not infiltrated by CD3+ T cells^106^. Cytology of tumors was performed at specified timepoints. Cellular specimens were collected using a 22-gauge needle attached to a 5 mL syringe. Following aspiration, samples were smeared onto a glass slide for subsequent cytochemical staining. Cytology slides were then evaluated by a board-certified veterinary pathologist. IHC staining and cytology samples were assessed on an Olympus BX45 microscope using a high-power 10x microscope objective. Digital images were captured used an Olympus DP28 digital camera and processed using Olympus cellSens Imaging Software (v4.2).

### Statistical analysis

Statistical analyses were conducted using Prism v10 (GraphPad). Power calculations were not conducted to predetermine sample size. The details of statistical analysis have been provided in the descriptions for figures.

## DATA AVAILABILITY

The data generated in this study are available within the article and its supplementary files. Nanostring expression data for canine tumor expression in dogs progressing after completion of RT with IL-2 and IL-12 therapy has been made publicly available in Gene Expression Omnibus (GEO) at GSE253243.

## ACKNOWLEDGEMENTS

We gratefully thank all of our pet dog owners for their consent and willingness to participate in this investigational trial. This study was supported by National Cancer Institute grant R01CA271243 (T.M.F. and K.D.W) and National Institute of Biomedical Imaging and Bioengineering grant R01EB031082 (K.D.W). A.S. was supported by National Science Foundation Graduate Research Fellowship Program. The authors would like to acknowledge Dr. Amy Schnelle and the Tumor Engineering and Phenotyping Shared Resource (TEP) at the Cancer Center at Illinois for assistance with histology and immunohistochemistry analysis.

## AUTHOR CONTRIBUTIONS

JAS, MMPB, AS, NM, KDW and TMF designed research; JAS, MMPB, AS, NM, EF, JH, KS, RK, KLB, and TMF performed research; JAS, MMPB, AS, NM, EF, KS, KLB, KDW, and TMF analyzed data; and JAS, MMPB, EF, TMF, and KDW wrote the paper.

## ETHICS DECLARATIONS - COMPETING INTERESTS

NM and KDW are named as inventors in a patent application filed by the Massachusetts Institute of Technology related to the data presented in this work (US20200102370A1). NM is an advisor to and KDW holds equity in Cullinan Oncology, which has licensed rights to the intellectual property mentioned above.

