## Extended Data File for "Tumor-localized interleukin-2 and interleukin-12 combine with radiation therapy to safely potentiate regression of advanced malignant melanoma in pet dogs"


Extended Data Figure 1: Additional patient characteristics and CT tumor response

Extended Data Figure 2: Local RT causes transient lymphodepletion in murine melanomas

Extended Data Figure 3: Kaplan-Meier survival plots of dogs treated for malignant melanoma

Extended Data Figure 4: IL-2/IL-12 cytokine dose does not appear to correlate with tumor response or survival

Extended Data Figure 5: Incidence of treatment-related adverse events

Extended Data Figure 6: IFN-γ and IL-10 response to cytokine-only or RT-only treatment

Extended Data Figure 7: Characterization of anti-drug antibody responses in treated dogs

Extended Data Figure 8: IHC analysis of delayed responder shows absence of melanoma

Extended Data Figure 9: IHC analysis of CD3 infiltration in progressor dogs

Extended Data Figure 10: Combined tumor and metastatic LN treatment response

| | At Presentation | | | | Primary Tumor Volume (in $cm^3$, by CT) | | |
|---|---|---|---|---|---|---|---|
| | Tumor Length (mm) | LN Disease | Metastasis | WHO Stage | Day 0 | Day 28 | Day 84 |
| **1x Dose Level Cohort** | | | | | | | |
| "Cricket" | 29 | Yes | Lung | IV | 4.7 | 2.4 | 3.5 |
| "Nala" | 30 | Yes | No | III | 7.5 | 6.5 | 6.1 |
| "Dezzi" | 26 | No | No | II | 11.6 | 2.8 | 0.1 |
| **2x Dose Level Cohort** | | | | | | | |
| "Maverick" | 31 | Yes | No | III | 3.5 | 0.3 | 0.0001 |
| "Max" | 28 | No | No | II | 7.6 | 0.5 | 0.7 |
| "Samba" | 8 | No | No | I | 0.5 | 0 | 0 |
| "Peanut" | 42 | Yes | No | III | 10.2 | 6.2 | 10.9 |
| "Izabella" | 27 | No | No | II | 5.9 | 0.3 | 0.4 |
| "Emily" | 30 | Yes | Skin | IV | 16.3 | 3.3 | *(euthanized)* |
| **3.3x Dose Level Cohort** | | | | | | | |
| "Marley" | not collected | Yes | No | III | 18.6 | 7.8 | 21.7 |
| "Komen" | not collected | Yes | No | III | 6.8 | 1.8 | 0.4 |
| "Candy" | 36 | Yes | Lung and Liver | IV | 9 | 0.2 | 0.5 |
| "Leahy" | 32 | No | No | II | 2.7 | 1.6 | 1.1 |
| **5x Dose Level Cohort** | | | | | | | |
| "Hank" | 65 | Yes | Lung | IV | 43.4 | 50.8 | *(euthanized)* |
| "Dexter" | 20 | Yes | Lung | IV | 3 | 0.1 | 0.0001 |

**Extended Data Figure 1: Additional patient characteristics and CT response data.** Ten of fifteen dogs presented with stage III or greater tumors at trial enrollment, with corresponding disease observed at tumor-draining lymph nodes or distal metastases. Radiologic response to treatment was measured using CT, with many patients displaying rapid and robust decreases in primary tumor volume.

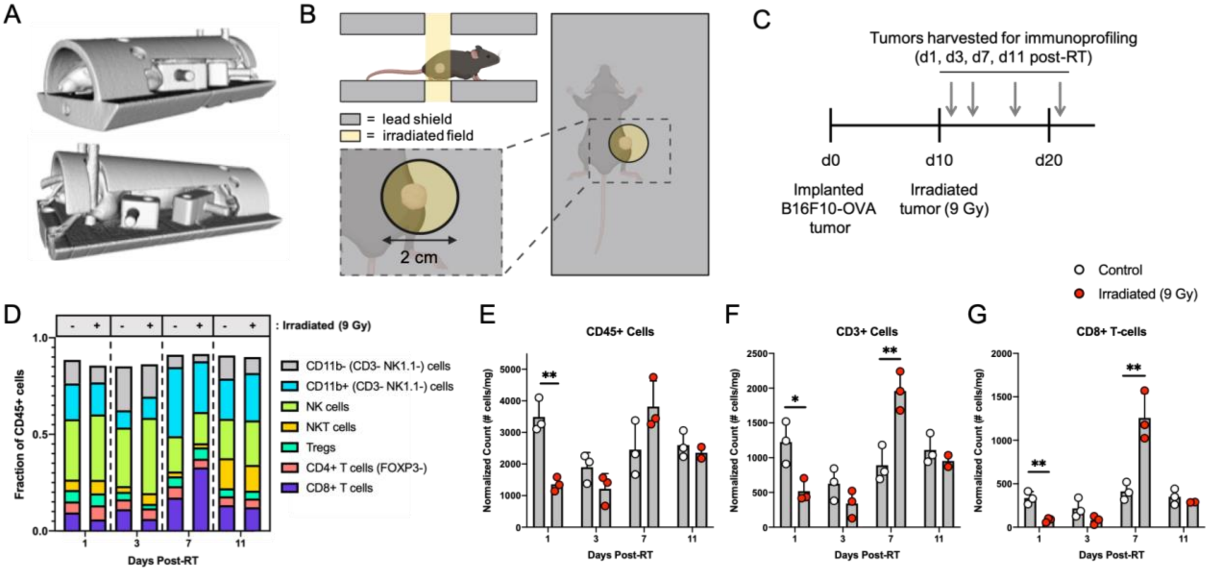


**Extended Data Figure 2: Local RT causes transient lymphodepletion in murine melanomas.** (A) 3D renderings of a mouse in the restrainer. The mouse is held in position by a pair of adjustable thorax and abdominopelvic blocks, and two sets of hind limb pins. (B) Schematic of local irradiation setup. The restrained mouse is placed between two lead shields (grey) and positioned to center the tumor within a 2 cm diameter aperture, which defines the irradiated field (yellow). A 9 Gy dose of radiation is delivered using a cesium-137 gamma irradiator. (C) Schema for immunophenotyping of tumors 1, 3, 7, and 11 days after irradiation. (D) Relative quantities of intratumoral immune populations (shown as fraction of CD45+ cells) following irradiation or control treatment, measured by flow cytometry. (E) Densities of intratumoral CD45+ cells (F) CD3+ cells (G) and CD8+ T cells following irradiation (red) or control (white) treatment (n = 3/group, mean + S.D.).

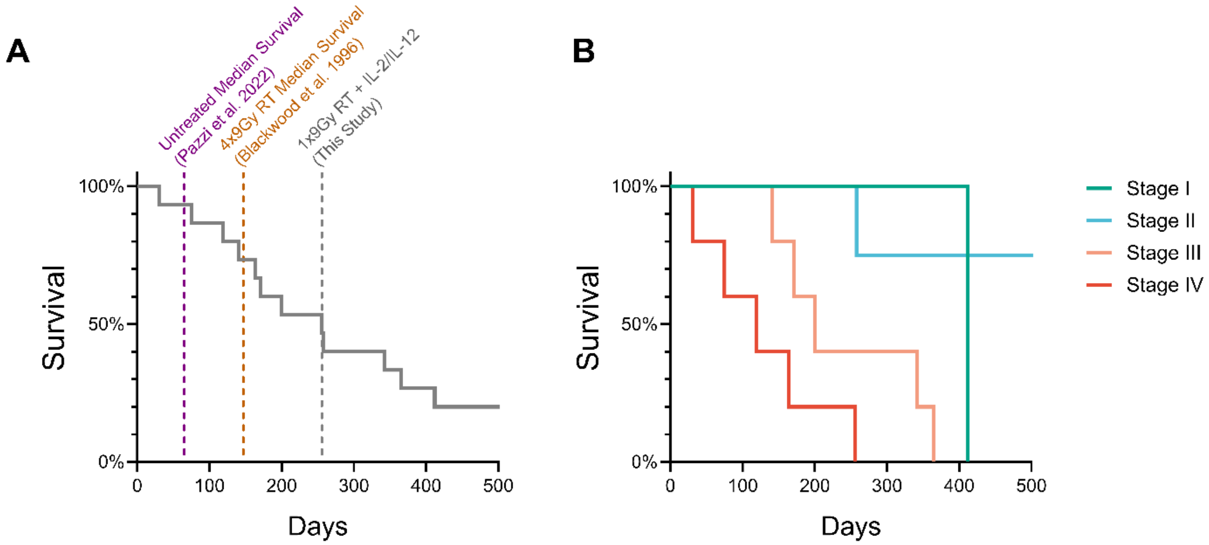


**Extended Data Figure 3: Kaplan-Meier survival plots of dogs treated for malignant melanoma.** (A) Kaplan-Meier plot showing survival time of all dogs treated with single 9 Gy dose of radiation therapy (RT) with up to six doses of intratumoral IL-2 and IL-12 cytokines, regardless of tumor stage and cytokine dose level. Dotted lines indicate the median survival times reported for untreated (65 days; purple) malignant melanoma and 4x9Gy RT treatment (147 days; orange) in dogs. Median survival for this study was 256 days across all treatment cohorts and tumor stages. (B) Kaplan-Meier plot showing survival time of dogs broken down by tumor stage at time of enrollment.

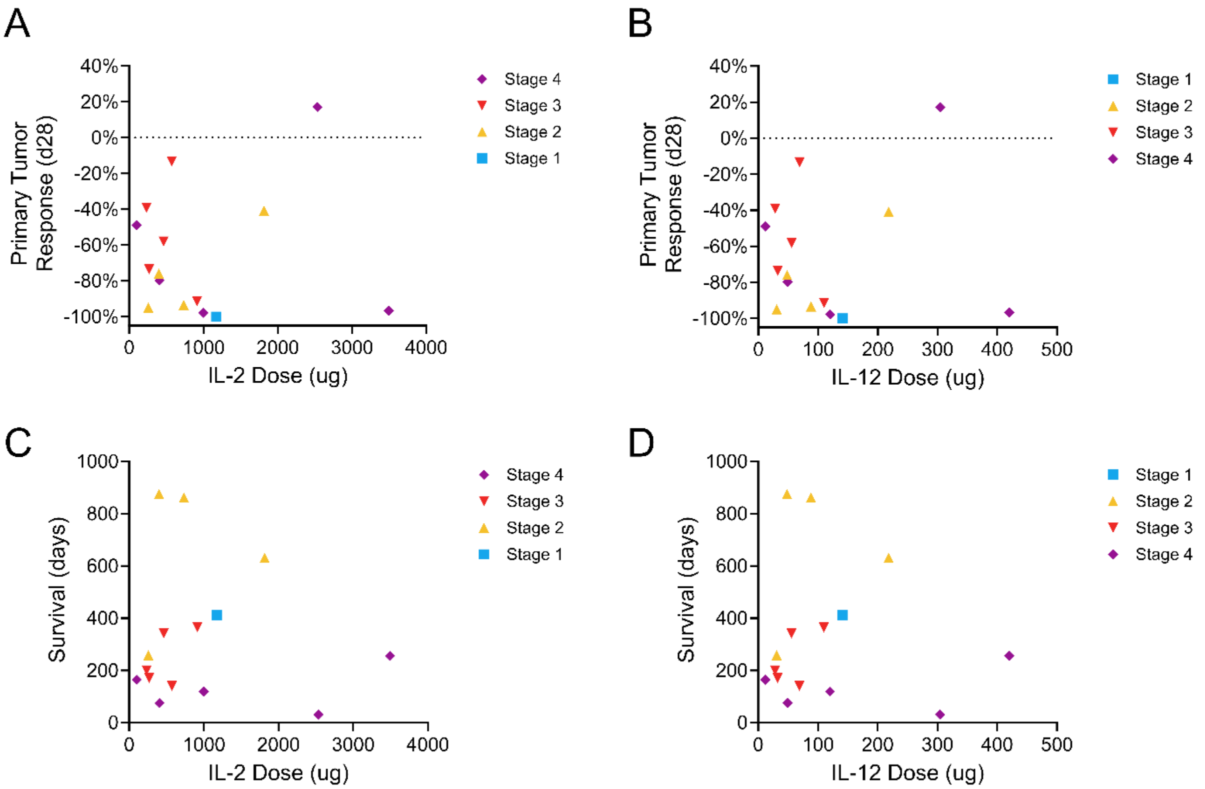


**Extended Data Figure 4: IL-2/IL-12 cytokine dose does not appear to correlate with tumor response or survival.** (A, B) Primary tumor responses measured at day 28 via CT are not improved at higher doses of IL-2 or IL-12 cytokine. (C, D) Similarly, overall survival appears to correlate more favorably with WHO tumor stage than IL-2 or IL-12 cytokine dose.

| | Cohort 1x (n= 3) | | | | | Cohort 2x (n= 6) | | | | | Cohort 3.3x (n= 4) | | | | | Cohort 5x (n= 2) | | | | | Total (n=15) |
|---|---|---|---|---|---|---|---|---|---|---|---|---|---|---|---|---|---|---|---|---|---|
| **Event** | Any grade | Grade 1 | Grade 2 | Grade 3 | Grade 4 | Any grade | Grade 1 | Grade 2 | Grade 3 | Grade 4 | Any grade | Grade 1 | Grade 2 | Grade 3 | Grade 4 | Any grade | Grade 1 | Grade 2 | Grade 3 | Grade 4 | |
| **Any event** | 40 | 33 (82.5%) | 7 (17.5%) | 0 | 0 | 168 | 93 (55.4%) | 54 (32.1%) | 18 (10.7%) | 3 (1.8%) | 122 | 84 (68.8%) | 28 (23%) | 10 (8.2%) | 0 | 84 | 36 (42.8%) | 31 (37%) | 15 (17.8%) | 2 (2.4%) | Total= 414 (100%) |
| **Blood/bone marrow** | | | | | | | | | | | | | | | | | | | | | |
| Hemoglobinemia | 5 | 5 (12.5%) | 0 | 0 | 0 | 16 | 13 (7.7%) | 2 (1.2%) | 1 (0.6%) | 0 | 14 | 11 (9%) | 3 (2.5%) | 0 | 0 | 8 | 7 (8.3%) | 1 (1.2%) | 0 | 0 | 43 (10.4%) |
| Thrombocytopenia | 4 | 4 (10%) | 0 | 0 | 0 | 12 | 10 (5.9%) | 2 (1.2%) | 0 | 0 | 11 | 8 (6.5%) | 3 (2.5%) | 0 | 0 | 6 | 2 (2.4%) | 3 (3.6%) | 0 | 1 (1.2%) | 33 (8%) |
| Neutropenia | 1 | 1 (2.5%) | 0 | 0 | 0 | 0 | 0 | 0 | 0 | 0 | 0 | 0 | 0 | 0 | 0 | 0 | 0 | 0 | 0 | 0 | 1 (0.2%) |
| Lymphocytosis | 0 | 0 | 0 | 0 | 0 | 1 | 0 | 1 (0.6%) | 0 | 0 | 0 | 0 | 0 | 0 | 0 | 0 | 0 | 0 | 0 | 0 | 1 (0.2%) |
| **Constitutional clinical signs** | | | | | | | | | | | | | | | | | | | | | |
| Fever | 4 | 1 (2.5%) | 3 (7.5%) | 0 | 0 | 11 | 6 (3.6%) | 5 (3%) | 0 | 0 | 6 | 1 (0.8%) | 3 (2.5%) | 2 (1.6%) | 0 | 4 | 0 | 3 (3.6%) | 1 (1.2%) | 0 | 25 (6%) |
| Lethargy/fatigue | 6 | 5 (12.5%) | 1 (2.5%) | 0 | 0 | 8 | 6 (3.6%) | 2 (1.2%) | 0 | 0 | 14 | 10 (8.2%) | 3 (2.5%) | 1 (0.8%) | 0 | 5 | 1 (1.2%) | 3 (3.6%) | 1 (1.2%) | 0 | 33 (8%) |
| **Dermatologic/Skin** | | | | | | | | | | | | | | | | | | | | | |
| Edema (injected region) | 1 | 0 | 1 (2.5%) | 0 | 0 | 3 | 3 (1.8%) | 0 | 0 | 0 | 4 | 3 (2.5%) | 1 (0.8%) | 0 | 0 | 2 | 2 (2.4%) | 0 | 0 | 0 | 10 (2.4%) |
| **Gastrointestinal** | | | | | | | | | | | | | | | | | | | | | |
| Diarrhea | 3 | 3 (7.5%) | 0 | 0 | 0 | 9 | 5 (3%) | 3 (1.8%) | 1 (0.6%) | 0 | 2 | 2 (1.6%) | 0 | 0 | 0 | 3 | 1 (1.2%) | 2 (2.4%) | 0 | 0 | 17 (4.1%) |
| Anorexia | 4 | 4 (10%) | 0 | 0 | 0 | 9 | 8 (4.8%) | 1 (0.6%) | 0 | 0 | 11 | 9 (7.4%) | 1 (0.8%) | 1 (0.8%) | 0 | 6 | 3 (3.6%) | 1 (1.2%) | 2 (2.4%) | 0 | 30 (7.2%) |
| Vomiting | 0 | 0 | 0 | 0 | 0 | 0 | 0 | 0 | 0 | 0 | 2 | 2 (1.6%) | 0 | 0 | 0 | 2 | 2 (2.4%) | 0 | 0 | 0 | 4 (1%) |
| Dehydration | 0 | 0 | 0 | 0 | 0 | 0 | 0 | 0 | 0 | 0 | 1 | 0 | 1 (0.8%) | 0 | 0 | 0 | 0 | 0 | 0 | 0 | 1 (0.2%) |
| **Metabolic** | | | | | | | | | | | | | | | | | | | | | |
| Increased ALT | 7 | 6 (15%) | 1 (2.5%) | 0 | 0 | 43 | 16 (9.5%) | 22 (13%) | 2 (1.2%) | 3 (1.8%) | 17 | 10 (8.2%) | 4 (3.3%) | 3 (2.5%) | 0 | 17 | 2 (2.4%) | 10 (11.9%) | 4 (4.7%) | 1 (1.2%) | 84 (20.3%) |
| Increased ALP | 4 | 4 (10%) | 0 | 0 | 0 | 33 | 8 (4.8%) | 11 (6.5%) | 14 (8.3%) | 0 | 28 | 19 (15.6%) | 6 (4.9%) | 3 (2.5%) | 0 | 16 | 6 (7.1%) | 4 (4.7%) | 6 (7.1%) | 0 | 81 (19.6%) |
| Increased CPK | 0 | 0 | 0 | 0 | 0 | 7 | 6 (3.6%) | 1 (0.6%) | 0 | 0 | 3 | 3 (2.5%) | 0 | 0 | 0 | 4 | 4 (4.7%) | 0 | 0 | 0 | 14 (3.4%) |
| Hypoalbuminemia | 0 | 0 | 0 | 0 | 0 | 15 | 12 (7.1%) | 3 (1.8%) | 0 | 0 | 3 | 2 (1.6%) | 1 (0.8%) | 0 | 0 | 8 | 5 (5.9%) | 3 (3.6%) | 0 | 0 | 26 (6.3%) |
| Increased Bilirubin | 0 | 0 | 0 | 0 | 0 | 1 | 0 | 1 (0.6%) | 0 | 0 | 4 | 3 (2.5%) | 1 (0.8%) | 0 | 0 | 3 | 1 (1.2%) | 1 (1.2%) | 1 (1.2%) | 0 | 8 (2%) |
| **Pain** | | | | | | | | | | | | | | | | | | | | | |
| Pain (injected region) | 1 | 0 | 1 (2.5%) | 0 | 0 | 0 | 0 | 0 | 0 | 0 | 2 | 1 (0.8%) | 1 (0.8%) | 0 | 0 | 0 | 0 | 0 | 0 | 0 | 3 (0.7%) |

**Extended Data Figure 5: Incidence of treatment-related adverse events.**

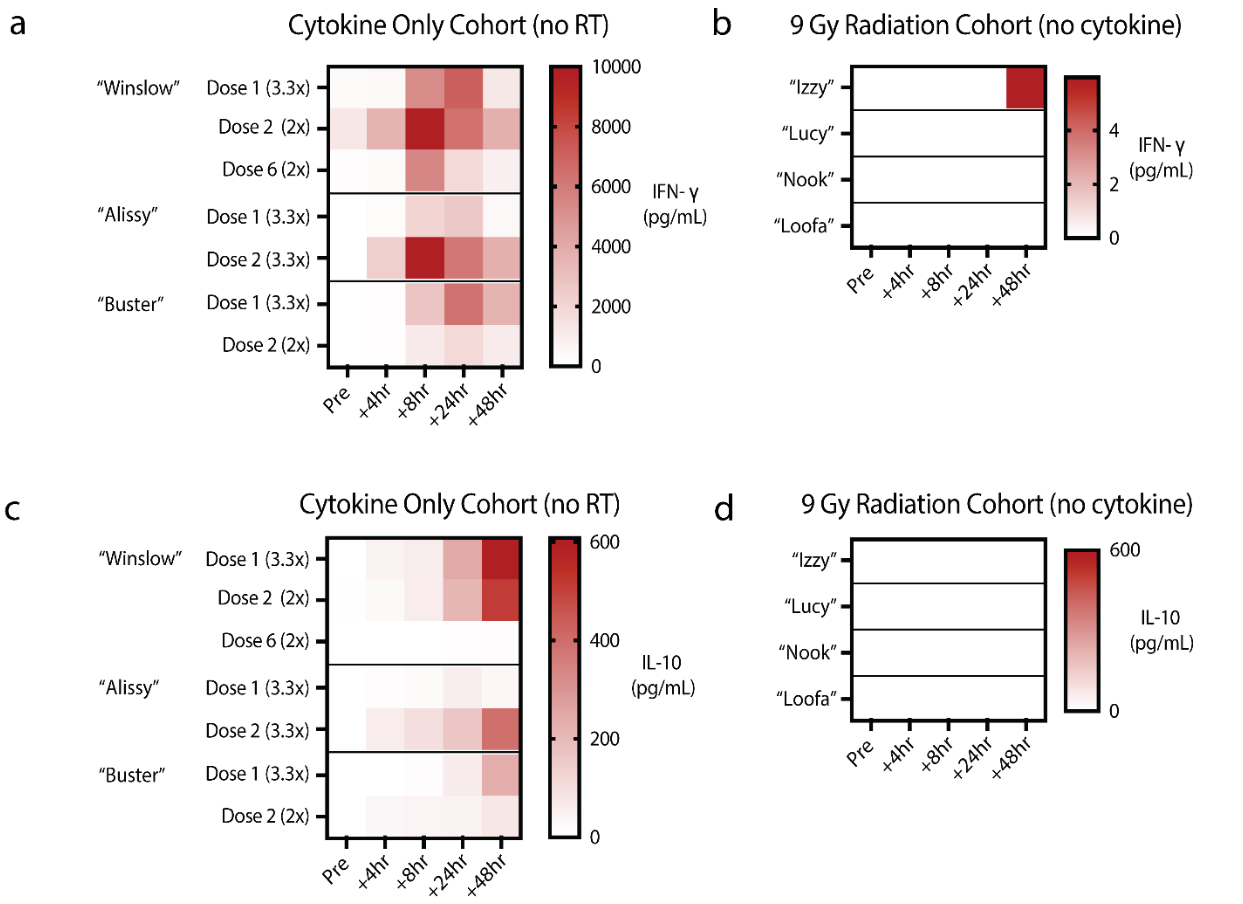


**Extended Data Figure 6: IFN-γ and IL-10 response to cytokine-only or RT-only treatment.** (a, b) The quantity of gamma interferon in patient serum was measured via ELISA at intervals following dosing of IL-2/IL-12 cytokines only (a) or 9 Gy radiation only (b). The increase in serum gamma interferon is only observed following intratumoral cytokine treatment. (c,d) Similarly, the amount of serum IL-10 was measured via ELISA following the same treatments. A delayed increase in IL-10 is only observed in the serum of dogs receiving intratumoral cytokine therapy (c) and was not detected in the serum of dogs receiving only a 9 Gy dose of radiation (d). This suggests that the systemic changes in cytokines/chemokines observed in Figure 2 are driven by the actions of the cytokine portion of the combination therapy, not the radiation dose alone.

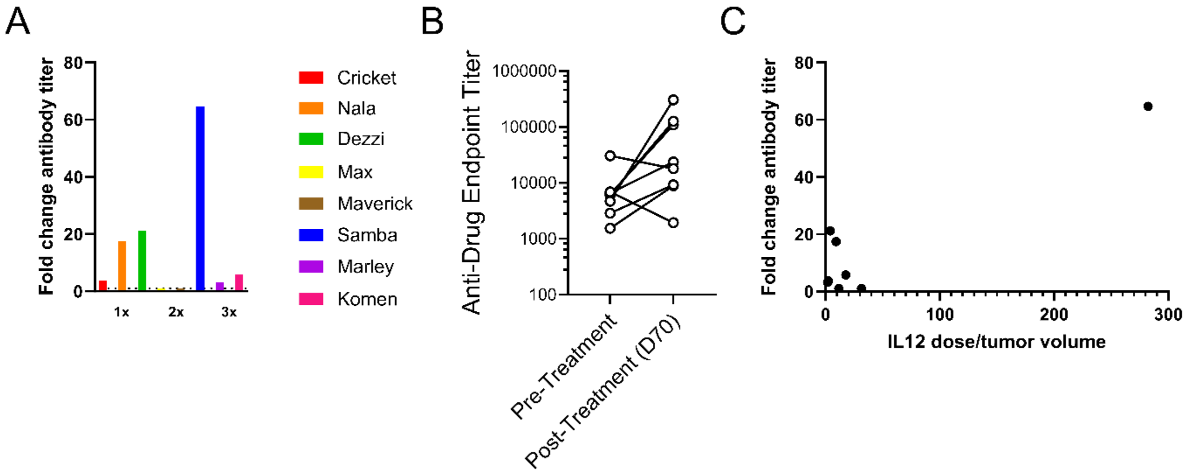


**Extended Data Figure 7: Characterization of anti-drug antibody responses in treated dogs.** (a) ELISA to characterize anti-drug antibodies against cLAIR-CSA-cIL2/cIL12-CSA-cLAIR was performed and fold-change in antibody titer was calculated using pre-treatment serum. (b) Endpoint titer values before and after receiving treatment was calculated as the reciprocal dilution at which 2x sample baseline was observed. (c) The ratio of dosed drug (ug) to tumor volume ($cm^3$) suggests that at doses exceeding the capacity of tumor retention through collagen binding, there may be an increased potential to raise anti-drug antibodies.

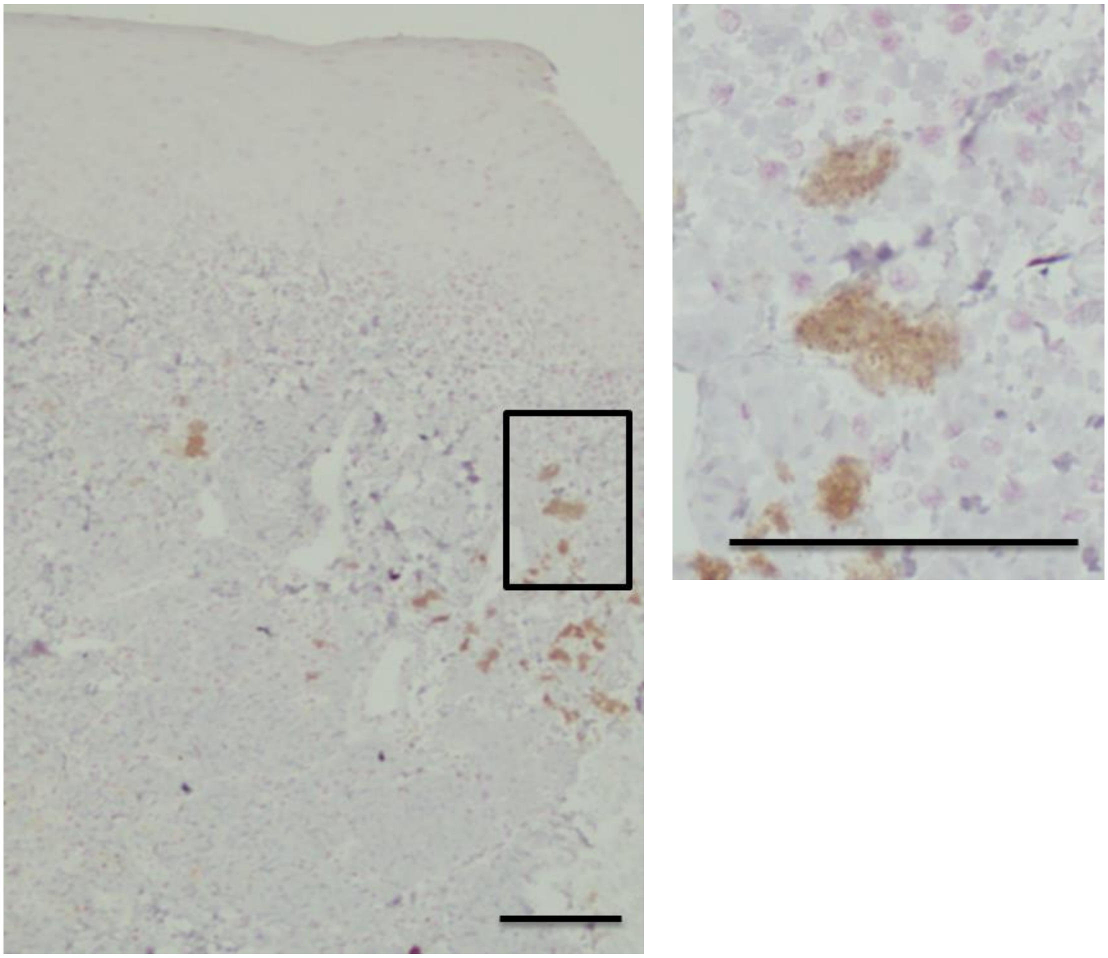

**Extended Data Figure 8: IHC analysis of delayed responder shows absence of melanoma.** No positive IHC signal is observed for Melan-A (brown) to indicate residual melanoma in the gingival tissue of the delayed responder. Brown pigment consistent with cytoplasmic accumulation of melanin is present within scattered melanophages (inset). Melan-A IHC with hematoxylin counterstain, scale bar 100µm.

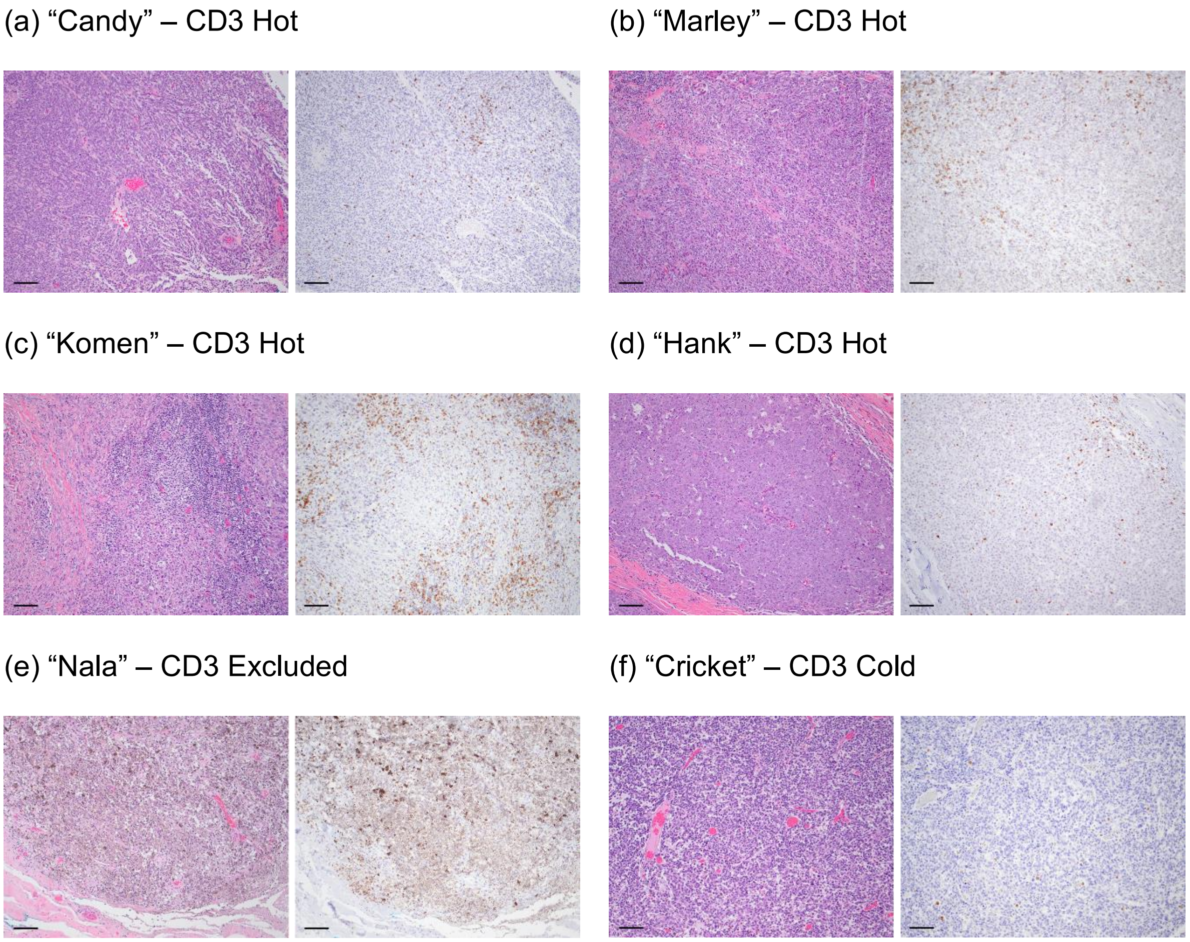


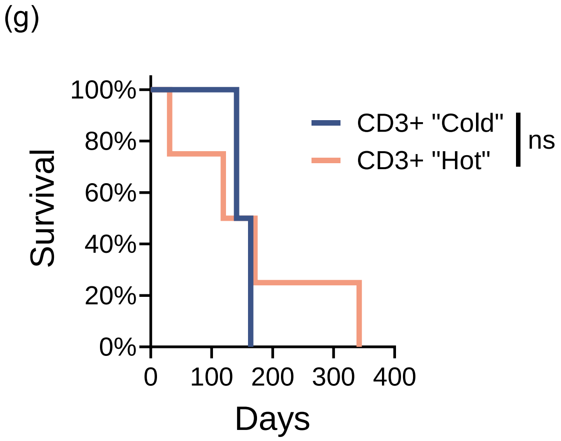


**Extended Data Figure 9: IHC analysis of CD3 infiltration in progressor dogs.** (a-f) For each dog, left image: H&E stained sample of primary tumor following patient euthanasia; right image: CD3 IHC staining for T cell infiltration at the same timepoint. Scale bar: 100um. (g) Overall survival of patients was not predicted by CD3 infiltration immunotype (hot vs. cold/excluded) using log-rank (Mantel-Cox) test. Ns: not significant.

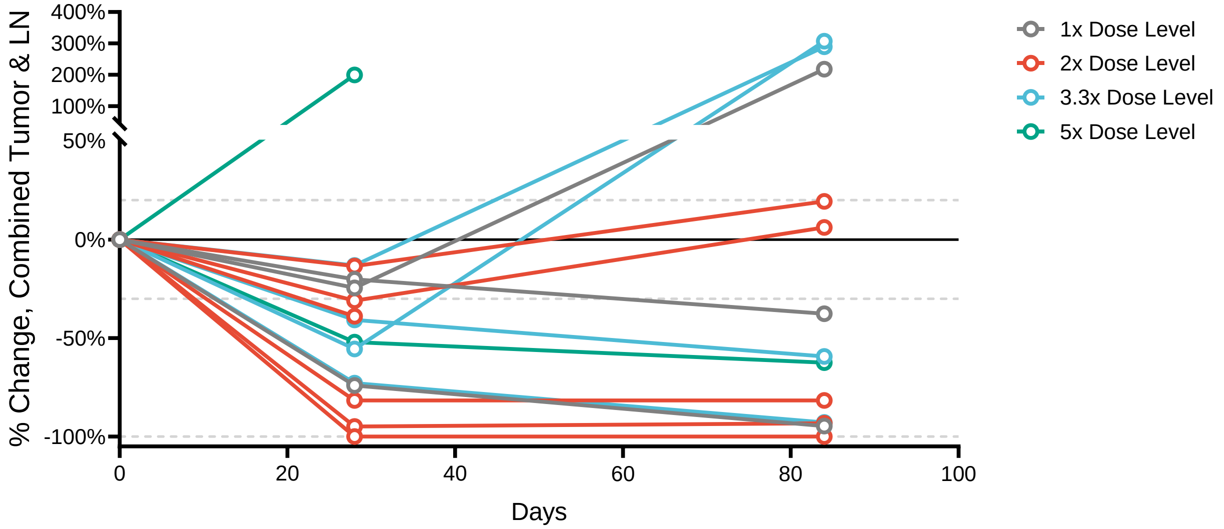


**Extended Data Figure 10: Combined tumor and metastatic LN treatment response.** CT measurements from primary tumor and diseased LN were combined to assess overall treatment response. Two dogs were euthanized prior to day 84 CT measurement due to progression of brain/CNS metastasis (not imaged via CT) or lung metastasis (not measured after CT). Dotted lines depict RECIST criteria for stable disease and partial response to treatment.
